# Sensitive periods for the effect of childhood adversity on DNA methylation: Results from a prospective, longitudinal study

**DOI:** 10.1101/271122

**Authors:** Erin C. Dunn, Thomas W. Soare, Andrew J. Simpkin, Matthew J. Suderman, Yiwen Zhu, Torsten Klengel, Andrew D.A.C. Smith, Kerry Ressler, Caroline L. Relton

**Author notes:** **Corresponding Author**: Erin C. Dunn, ScD, MPH, Psychiatric and Neurodevelopmental Genetics Unit, Center for Human Genetic Research, Massachusetts General Hospital, 185 Cambridge Street, Simches Research Building 6th Floor, Boston, MA 02114; email: edunn[at]mgh[dot]harvard[dot]edu.; website: www.thedunnlab.com.

## Abstract

**Background:** Exposure to “early life” adversity is known to predict DNA methylation (DNAm) patterns that may be related to prolonged psychiatric risk. However, few studies have investigated whether adversity has time-dependent effects based on the age at exposure.

**Methods:** Using a two-stage structured life course modeling approach (SLCMA), we tested the hypothesis that there are sensitive periods when adversity induced greater DNAm changes. We tested this hypothesis in relation to two alternative explanations: an accumulation hypothesis, in which the effect of adversity on DNAm increases with the number of occasions exposed, regardless of timing, and a recency model, in which the effect of adversity is stronger for more proximal events. Data came from the Accessible Resource for Integrated Epigenomics Studies (ARIES), a subsample of mother-child pairs from the Avon Longitudinal Study of Parents and Children (ALSPAC; n=670-776).

**Results:** After covariate adjustment and multiple testing correction, we identified 40 CpG sites that were differentially methylated at age 7 following exposure to adversity. Most loci (n=32) were predicted by the timing of adversity, namely exposures during infancy. Neither the accumulation nor recency of the adversity explained considerable variability in DNAm. A standard EWAS of lifetime exposure (vs. no exposure) failed to detect these associations.

**Conclusions:** The developmental timing of adversity explains more variability in DNAm than the accumulation or recency of exposure. Infancy appears to be a sensitive period when exposure to adversity predicts differential DNAm patterns. Classification of individuals as exposed vs. unexposed to “early life” adversity may dilute observed effects.

## Introduction

Exposure to childhood adversity, including poverty (1), abuse (2, 3), family disruption or dysfunction (4, 5), and other stressors (6, 7), is a common and potent determinant of mental health across the lifespan, increasing risk of childhood– and adult-onset psychiatric disorders by at least two-fold (8–10). Although the biological mechanisms explaining this relationship are poorly understood, accumulating evidence suggests adversity may become programmed molecularly, leaving behind biological memories that persistently alter genome function and increase susceptibility to mental disorders. Indeed, dozens of candidate gene and epigenome-wide association studies (EWAS) in both animals and humans have shown that early life adversity is associated with persistent alterations in the epigenome (11–15), including changes in DNA methylation (DNAm), which is the most studied epigenetic mechanism involving the addition of methyl groups to cytosines in the DNA sequence (16, 17). These differential DNAm sites can robustly alter specific gene expression, providing an epigenetic mechanism by which gene by environment interactions directly affect biological responses (18).

Recent evidence, particularly from animal studies, suggests that epigenetic programming may be developmentally time-sensitive and that there may be sensitive periods (19, 20) when adversity exposure is more likely to induce DNAm changes. For instance, rodent experiments have demonstrated the existence of sensitive periods for different aspects of epigenetic regulation - from embryonic reprogramming to postnatal exposure leading to differences in epigenetic outcomes and gene expression (21–25). Recent work in nonhuman primates also suggests that there are differential effects on DNAm profiles based on whether adversity exposure, including maternal separation, occurred at birth versus later stages of development (26). Yet, few human studies, whether candidate gene (16, 27–29) or EWAS (30–32), have examined the time-dependent effects of psychosocial adversity on DNAm; nearly all human epigenetic studies have instead focused on the presence versus absence of exposure to “early life” adversity. Thus, it is unknown whether there are age stages when adversity exposure differentially affects DNAm, and when children are therefore more vulnerable and prevention efforts could be most efficacious.

This study aimed to address this limitation by using data from a prospective, longitudinal birth cohort of young children to test the hypothesis that there are sensitive periods associated with DNAm alterations following adversity exposure. To test this hypothesis, we utilized a two-stage Structured Life Course Modeling Approach (SLCMA) (33, 34) to examine the effect of repeated exposure to seven types of childhood adversities across three developmental periods (in infancy, before age 3; preschool, ages 3-5; and middle childhood, ages 6-7) on DNAm profiles measured at age 7. Recognizing that alternative conceptual models have been proposed to explain the effects of adversity, we also used the SLCMA to determine whether the sensitive period model explained more variability in DNAm relative to two other theoretical models described in the life course epidemiology literature (35–37): (1) an accumulation model (38–40), in which the effect of adversity on DNAm increases with the number of occasions exposed, regardless of timing; and (2) a recency model (41), in which the effect of adversity on DNAm is stronger for more proximal events. Finally, to evaluate the potential advantage of the SLCMA relative to the standard EWAS approach, which would ignore the timing or frequency of adversity, we examined the number of epigenome-wide significant loci identified by each approach and evaluated their degree of overlap.

## Methods and Materials

### Sample and Procedures

Data came from the Avon Longitudinal Study of Parents and Children (ALSPAC), a population-based birth cohort (42–44). ALSPAC generated blood-based DNAm profiles at birth and age 7 as part of the Accessible Resource for Integrated Epigenomics Studies (ARIES), a subsample of 1,018 mother-child pairs from the ALSPAC (45). The ARIES mother-child pairs were randomly selected out of those with complete data across at least five waves of data collection (**Supplemental Materials)**.

### Measures

#### Exposure to Adversity

We examined the effect of seven adversities shown previously to associate with epigenetic marks (46–48): (a) caregiver physical or emotional abuse (49–52); (b) sexual or physical abuse (by anyone) (49–52); (c) maternal psychopathology (53, 54); (d) one adult in the household (55); (e) family instability (56, 57); (f) financial stress/poverty (58, 59); and (g) neighborhood disadvantage/poverty (60). Each adversity was measured via maternal report on at least four occasions at or before age 7 from a single item or psychometrically validated standardized measures. Specific time periods of assessment varied across adversity type (**Supplemental Methods**). For each type of adversity, we generated three sets of encoded variables (**Supplemental Materials**): (a) a set of variables indicating presence versus absence of the adversity at a specific developmental stage, to test the *sensitive period hypothesis*; (b) a single variable denoting the total number of time periods of exposure to a given adversity, to test the *accumulation hypothesis*; and (c) a single variable denoting the total number of developmental periods of exposure, with each exposure weighted by the age of the child during the measurement time period, to test the *recency hypothesis*; this recency variable upweighted more recent exposures, allowing us to determine whether more recent exposures were more impactful.

#### DNA Methylation

DNAm was measured at 485,000 CpG dinucleotide sites across the genome using the Illumina Infinium Human Methylation 450k BeadChip microarray, which captures DNAm variation at 99% of RefSeq genes (17 CpG sites per gene, on average). DNA for this assay was extracted from cord blood and peripheral blood leukocytes at age 7. DNA methylation wet laboratory procedures, preprocessing analyses, and quality control were performed at the University of Bristol (**Supplemental Materials** and (45)). DNAm levels are expressed as a ‘beta’ value (β-value), representing the proportion of cells methylated at each interrogated CpG site and ranges from 0 (no methylated dinucleotides observed) to 1 (all dinucleotides methylated).

Prior to analysis, raw methylation β-values, which are preferred over M-values due to their interpretability (61), were normalized (62) to remove or minimize the effects of variation due to technical artifacts. To adjust for DNAm variation due to cell type heterogeneity in peripheral and cord blood samples, we estimated cell counts from DNAm profiles (63) and regressed out these estimates from the normalized β-values. Additionally, to remove possible outliers, we winsorized the β-values at each CpG site, setting the bottom 5% and top 95% of values to the 5th and 95th quantile, respectively (64).

#### Covariates

To adjust for baseline socio-demographic differences in the cohort, all analyses additionally controlled for the following variables, measured at child birth (**Supplemental Methods**): child race/ethnicity; child birth weight; maternal age; singleton vs. multiple birth; number of previous pregnancies; sustained maternal smoking during pregnancy; and parent social class (65). Parent social class was included because it captures a more fixed aspect of social class, encompassing job industry and rank, as well as education, wealth, social status and other aspects of socioeconomic status (66), thus allowing us to control for potential confounding effects of the social class into which children are born. Inclusion of parent social class also did not preclude the examination of financial adversity, which is a more temporally dynamic aspect of socioeconomic status.

### Data Analysis

Our primary analyses involved comparing the three theoretical models using the SLCMA, which was originally developed by Mishra (67) and later extended by Smith (33, 34) to analyze repeated, binary exposure data across the life course. The major advantage of the SLCMA is that it provides an unbiased way to compare multiple competing theoretical models simultaneously and identify the most parsimonious explanation for the observed outcome variation. The SLCMA uses Least Angle Regression (LARS) (68) and an associated covariance test (69) to identify the single theoretical model (or potentially more than one model working in combination) that explains the most outcome variation (R^2^). Compared to other methods for structured life course analysis, LARS has greater statistical power (33) and does not over-inflate effect size estimates (68) or bias hypothesis tests (69). The SLCMA has been used in several life course epidemiology studies (70, 71), including studies of other birth cohorts (72, 73). The LARS procedure functions under the same assumptions as multiple linear regression.

In the first stage, we entered the set of encoded variables described previously into the LARS variable selection procedure (68). LARS identified the variable with the strongest association with the outcome, thus identifying whether the sensitive period, accumulation, or recency model was most supported by the data. Therefore, *for each CpG site*, seven unique LARS models were selected, corresponding to each type of adversity. For each selected model, we performed a covariance test of the null hypothesis that the variable selected is unassociated with the outcome. With respect to multiple testing, the covariance test p-values are adjusted for the number of variables included in the LARS procedure, controlling the type I error rate for each adversity type and CpG site. To adjust for confounding during the first stage, we regressed each encoded variable on the covariates and implemented LARS on the regression residuals (34).

In the second stage, the theoretical model shown in the first stage to best fit the observed data for a specific type of adversity was then carried forward to a multivariate regression framework, where measures of effect were estimated. Only models with a covariance test p-value <1×10^−7^, the standard Bonferroni correction threshold for epigenome-wide statistical significance, were included in the second stage. Positive effect estimates thus indicate elevated (hyper) methylation and negative effect estimates indicate decreased (hypo) methylation. The same covariates were also included in the second stage. We compared the distribution of theoretical models across the Bonferroni-significant CpG sites with an omnibus chi-squared test, which tested the null hypothesis that the theoretical models were likely to be represented among the significant results in proportion to the frequency in which they were tested.

To evaluate the loss or gain of information when using a simpler versus more complex analytic approach, we also performed seven EWASs (one for each type of adversity) to evaluate the association between lifetime exposure to adversity (coded as ever versus never exposed) and DNAm across all CpG sites. The EWAS results were then compared to the SLCMA to determine if the two approaches yielded similar or distinct conclusions regarding the number of significant loci detected.

We also performed sensitivity analyses to evaluate the fit of the LARS selection procedure, determine the degree of differential methylation present at birth, and control for genetic variation. We examined the biological significance of the findings by: (a) examining the correlation in methylation between blood and brain tissue for the top CpG sites using an online database (74); (b) investigating enrichment of regulatory elements annotated to false discovery rate (FDR)-significant CpG sites; (c) performing a functional clustering analysis of all Gene Ontology (GO) terms for genes annotated to FDR-significant sites in DAVID 6.8 (75); (d) assessing the selective constraint of these genes using the Exome Aggregation Consortium (ExAC) (76); and (e) searching the NHGRI-EBI GWAS Catalog (GRASP, (77)) for phenotypic associations with genes annotated to the top CpG sites.

## Results

### Sample Characteristics and Distribution of Exposure to Adversity

Demographic characteristics of the ARIES analytic sample are shown in **Table S1** for the total sample and among children exposed to any adversity (n=744, 76%, experienced at least one adversity at some point in their lifetime). Details on the prevalence and correlations of exposure across time are also reported in **Figures 1** **and S1 and Table S2**. Of note, differences in the prevalence of exposure across time are unlikely to affect model selection as all variables are automatically standardized by the LARS procedure.

**Figure 1.**
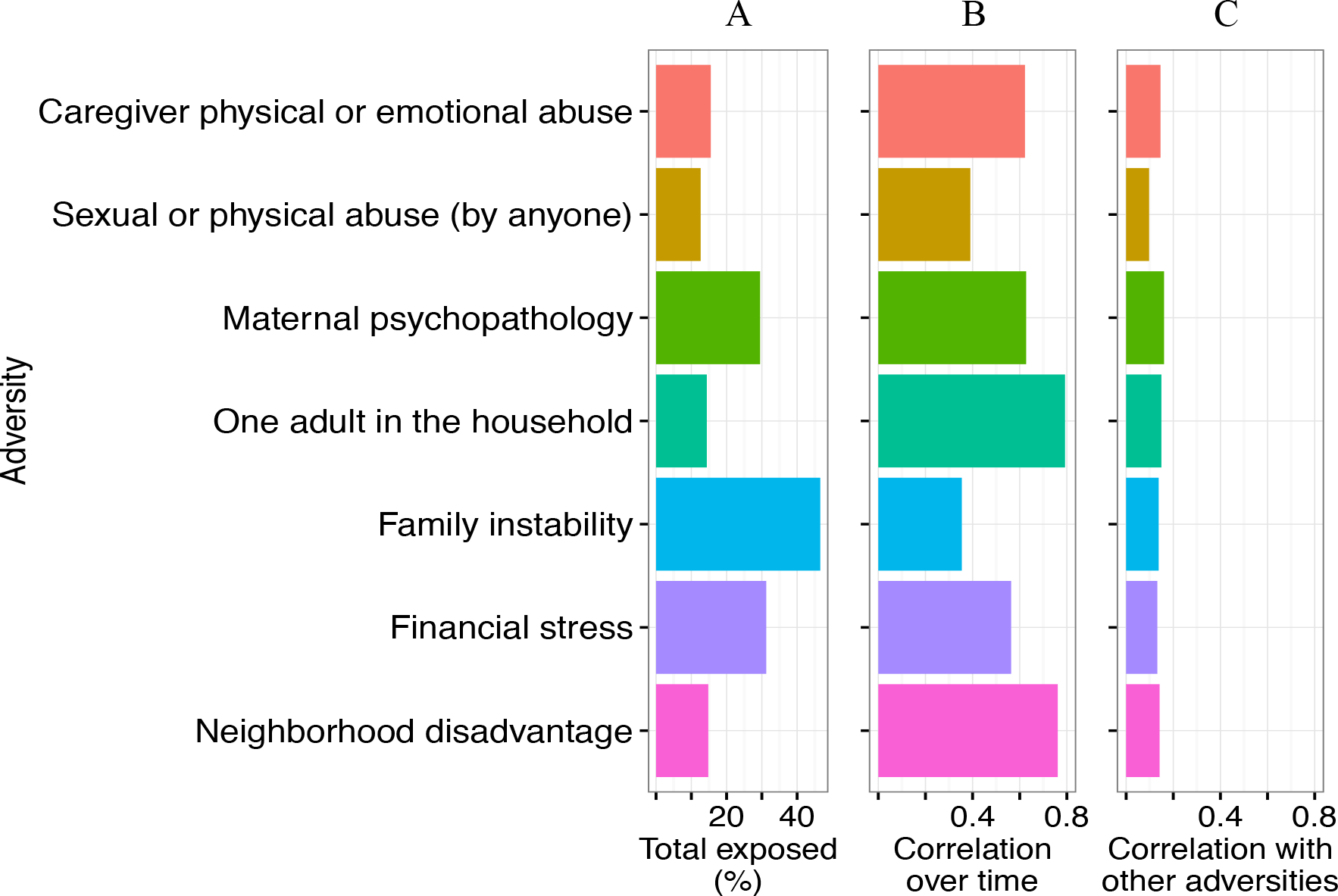
Exposure to adversity in the ARIES dataset. The figure displays the lifetime prevalence by age 7 of exposure to each adversity (labeled as *total exposed*), the average correlation between exposure to one type of adversity at one time point with exposure to that same adversity at a second time point (labeled as *correlation over time*), and the average correlation between exposure to one type of adversity and a second type of adversity (labeled as *correlation with other adversities*). **Panel A**: The lifetime prevalence of each adversity varied by type. The most commonly reported adversities were family instability (47%), maternal psychopathology (29%), and financial stress (27%). The remaining adversities were the least reported adversities, but still common: One adult in the household (12%), caregiver physical or emotional abuse (12%), sexual or physical abuse (by anyone; 13%), and neighborhood disadvantage (15%). **Panel B**: Among specific types of adversity, exposures tended to correlate over time, with neighboring time points being more related than distant time points. For instance, exposure to one adult in the household and neighborhood disadvantage were most strongly correlated over time (r=0.57-0.93 and r=0.70-0.88, respectively), whereas exposure to family instability (r=0.14-0.66) and sexual or physical abuse (r=0.23-0.65) were more weakly correlated across time. **Panel C**: The average correlation of exposure to the other adversities was modest across adversities, suggesting that we were capturing unique subtypes of adversity.

### Model Comparison and Effect Estimation

We identified 40 CpG sites (“top sites”) that were differentially methylated at age 7 following exposure to adversity (p<1×10^−7^, **Figure 2**). Methylation at most sites (n=38) was related to the developmental timing of exposure to adversity, especially adversity during infancy, meaning between birth and age 2 (**Figure 3a**). In fact, exposure to adversity during infancy explained variability at more CpG sites (23 in total) than expected, while the accumulation and recency models were associated with fewer CpG sites than expected (0 and 2 CpG sites, respectively; *χ*^2^=13.36, p=0.01).

**Figure 2.**
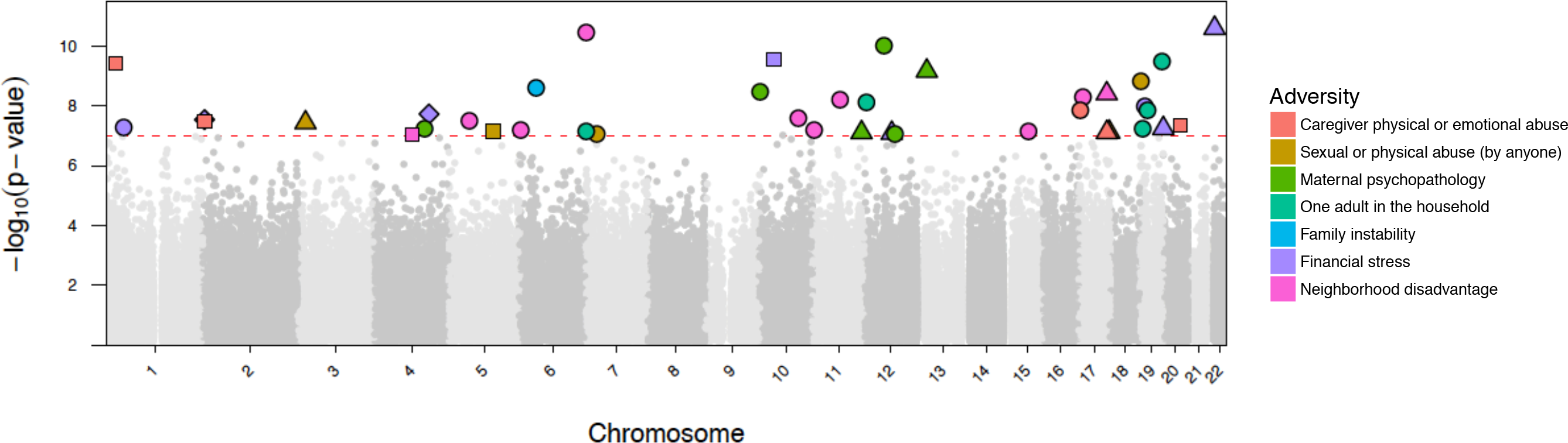
Manhattan plot displaying top CpG sites associated with exposure to adversity. In this Manhattan plot, the x-axis is the chromosomal position for each CpG site and the y-axis is the −log10 p-value for the association between exposure to adversity and DNAm values at each CpG site. The dashed line shows the epigenome-wide significance level, with each CpG site above the line representing a statistically significant association (p<1×10^−7^). The color of each CpG site refers to the type of adversity. The shape of each CpG site indicates the lifecourse model tested. The sensitive period hypotheses were encoded as *circle*: infancy, *triangle*: preschool, *square*: middle childhood. The recency hypothesis was encoded as a *diamond*. As shown, CpG sites significantly affected by exposure adversity were distributed throughout the genome. There was no obvious genomic spatial pattern by adversity type or timing of exposure.

**Figure 3.**
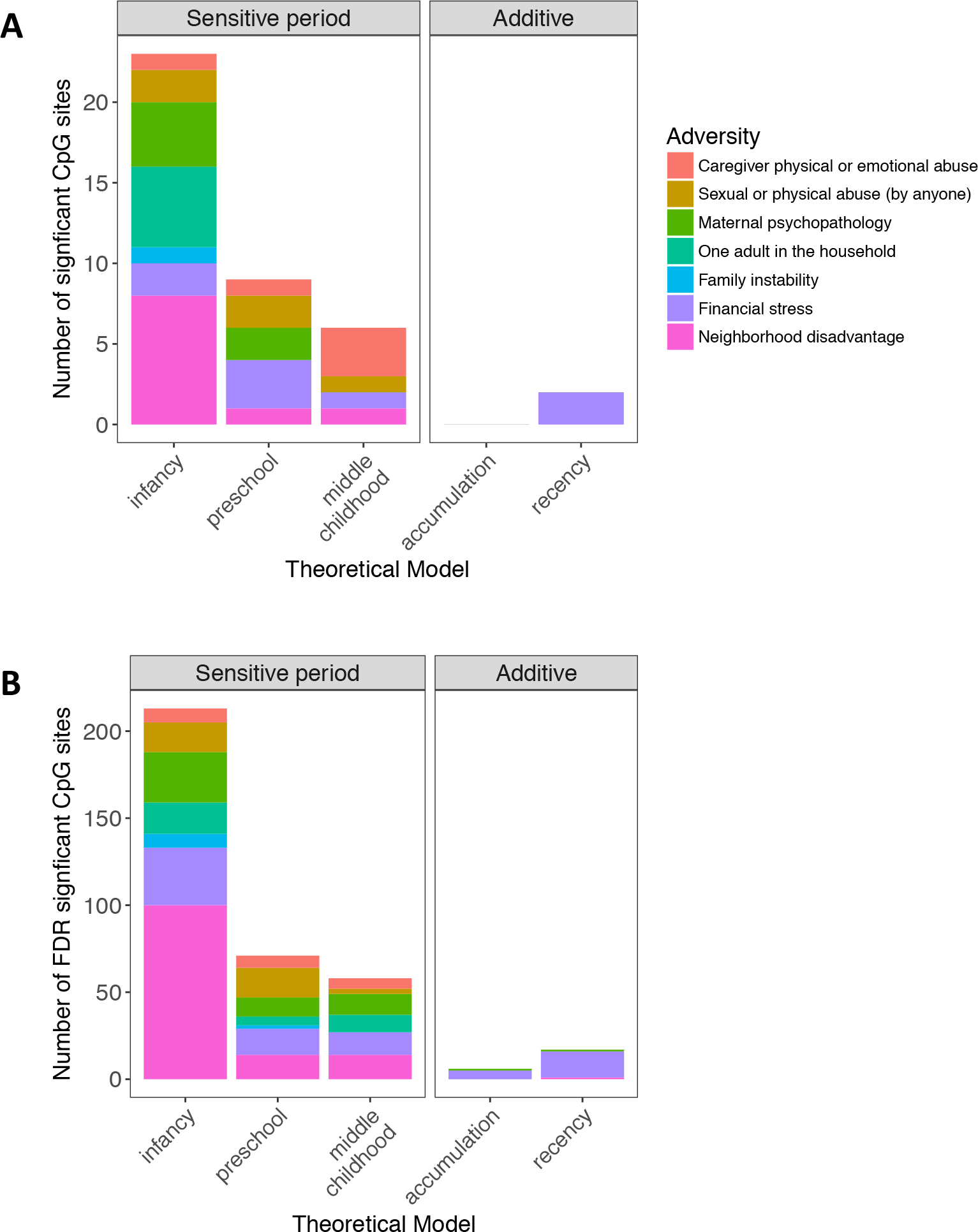
Frequency each lifecourse theoretical model was chosen for each type of adversity. Each plot displays the number of CpG sites for which adversity significantly predicted methylation, after controlling for covariates and correcting for multiple comparisons using **(a)** a Bonferroni threshold (p<1×10^−7^, n=40 sites) and **(b)** a False Discovery Rate (FDR) correction q < 0.05 (n=365 sites). The distribution of theoretical models chosen first by the LARS procedure for top CpG sites was significantly different than expected by chance, with exposure to adversity during sensitive periods, especially during infancy, more frequently predicting methylation.

As shown in **Table 1** and **Figure 3a**, neighborhood disadvantage was the type of adversity predicting the greatest number of genome-wide methylation differences (10 CpG sites), followed by financial stress (8 CpG sites) and maternal psychopathology (6 CpG sites). The remaining adversities were associated with differences at five CpG sites each, except for family instability, which was associated with differential methylation at a single CpG site.

**Table 1.**
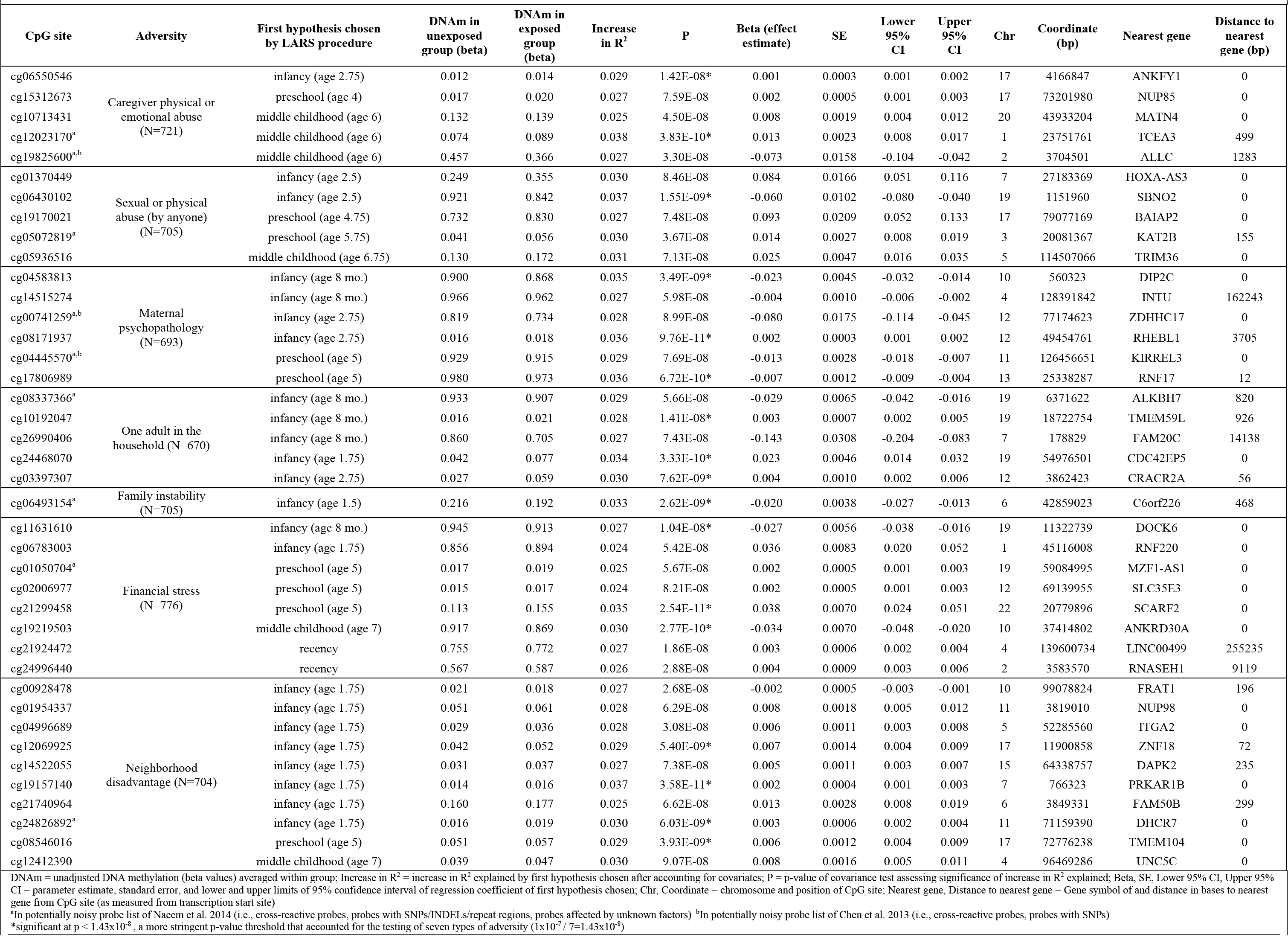
Results of the Structured Lifecourse Modeling Approach (SLCMA) in ARIES, with annotation to the closest gene, for the Bonferroni-significant CpG sites (p<1×10^−7^).

| CpG site | Adversity | First hypothesis chosen by LARS procedure | DNAm in unexposed group (beta) | DNAm in exposed group (beta) | Increase in $R^2$ | P | Beta (effect estimate) | SE | Lower 95% CI | Upper 95% CI | Chr | Coordinate (bp) | Nearest gene | Distance to nearest gene (bp) |
| --- | --- | --- | --- | --- | --- | --- | --- | --- | --- | --- | --- | --- | --- | --- |
| cg06550546 | Caregiver physical or emotional abuse (N=721) | infancy (age 2.75) | 0.012 | 0.014 | 0.029 | 1.42E-08* | 0.001 | 0.0003 | 0.001 | 0.002 | 17 | 4166847 | ANKFY1 | 0 |
| cg15312673 |  | preschool (age 4) | 0.017 | 0.020 | 0.027 | 7.59E-08 | 0.002 | 0.0005 | 0.001 | 0.003 | 17 | 73201980 | NUP85 | 0 |
| cg10713431 |  | middle childhood (age 6) | 0.132 | 0.139 | 0.025 | 4.50E-08 | 0.008 | 0.0019 | 0.004 | 0.012 | 20 | 43933204 | MATN4 | 0 |
| cg12023170<sup>a</sup> |  | middle childhood (age 6) | 0.074 | 0.089 | 0.038 | 3.83E-10* | 0.013 | 0.0023 | 0.008 | 0.017 | 1 | 23751761 | TCEA3 | 499 |
| cg19825600<sup>a,b</sup> |  | middle childhood (age 6) | 0.457 | 0.366 | 0.027 | 3.30E-08 | -0.073 | 0.0158 | -0.104 | -0.042 | 2 | 3704501 | ALLC | 1283 |
| cg01370449 | Sexual or physical abuse (by anyone) (N=705) | infancy (age 2.5) | 0.249 | 0.355 | 0.030 | 8.46E-08 | 0.084 | 0.0166 | 0.051 | 0.116 | 7 | 27183369 | HOXA-AS3 | 0 |
| cg06430102 |  | infancy (age 2.5) | 0.921 | 0.842 | 0.037 | 1.55E-09* | -0.060 | 0.0102 | -0.080 | -0.040 | 19 | 1151960 | SBNO2 | 0 |
| cg19170021 |  | preschool (age 4.75) | 0.732 | 0.830 | 0.027 | 7.48E-08 | 0.093 | 0.0209 | 0.052 | 0.133 | 17 | 79077169 | BAIAP2 | 0 |
| cg05072819<sup>a</sup> |  | preschool (age 5.75) | 0.041 | 0.056 | 0.030 | 3.67E-08 | 0.014 | 0.0027 | 0.008 | 0.019 | 3 | 20081367 | KAT2B | 155 |
| cg05936516 |  | middle childhood (age 6.75) | 0.130 | 0.172 | 0.031 | 7.13E-08 | 0.025 | 0.0047 | 0.016 | 0.035 | 5 | 114507066 | TRIM36 | 0 |
| cg04583813 | Maternal psychopathology (N=693) | infancy (age 8 mo.) | 0.900 | 0.868 | 0.035 | 3.49E-09* | -0.023 | 0.0045 | -0.032 | -0.014 | 10 | 560323 | DIP2C | 0 |
| cg14515274 |  | infancy (age 8 mo.) | 0.966 | 0.962 | 0.027 | 5.98E-08 | -0.004 | 0.0010 | -0.006 | -0.002 | 4 | 128391842 | INTU | 162243 |
| cg00741259<sup>a,b</sup> |  | infancy (age 2.75) | 0.819 | 0.734 | 0.028 | 8.99E-08 | -0.080 | 0.0175 | -0.114 | -0.045 | 12 | 77174623 | ZDHHC17 | 0 |
| cg08171937 |  | infancy (age 2.75) | 0.016 | 0.018 | 0.036 | 9.76E-11* | 0.002 | 0.0003 | 0.001 | 0.002 | 12 | 49454761 | RHEBL1 | 3705 |
| cg04445570<sup>a,b</sup> |  | preschool (age 5) | 0.929 | 0.915 | 0.029 | 7.69E-08 | -0.013 | 0.0028 | -0.018 | -0.007 | 11 | 126456651 | KIRREL3 | 0 |
| cg17806989 |  | preschool (age 5) | 0.980 | 0.973 | 0.036 | 6.72E-10* | -0.007 | 0.0012 | -0.009 | -0.004 | 13 | 25338287 | RNF17 | 12 |
| cg08337366<sup>a</sup> | One adult in the household (N=670) | infancy (age 8 mo.) | 0.933 | 0.907 | 0.029 | 5.66E-08 | -0.029 | 0.0065 | -0.042 | -0.016 | 19 | 6371622 | ALKBH7 | 820 |
| cg10192047 |  | infancy (age 8 mo.) | 0.016 | 0.021 | 0.028 | 1.41E-08* | 0.003 | 0.0007 | 0.002 | 0.005 | 19 | 18722754 | TMEM59L | 926 |
| cg26990406 |  | infancy (age 8 mo.) | 0.860 | 0.705 | 0.027 | 7.43E-08 | -0.143 | 0.0308 | -0.204 | -0.083 | 7 | 178829 | FAM20C | 14138 |
| cg24468070 |  | infancy (age 1.75) | 0.042 | 0.077 | 0.034 | 3.33E-10* | 0.023 | 0.0046 | 0.014 | 0.032 | 19 | 54976501 | CDC42EP5 | 0 |
| cg03397307 |  | infancy (age 2.75) | 0.027 | 0.059 | 0.030 | 7.62E-09* | 0.004 | 0.0010 | 0.002 | 0.006 | 12 | 3862423 | CRACR2A | 56 |
| cg06493154<sup>a</sup> | Family instability (N=705) | infancy (age 1.5) | 0.216 | 0.192 | 0.033 | 2.62E-09* | -0.020 | 0.0038 | -0.027 | -0.013 | 6 | 42859023 | C6orf226 | 468 |
| cg11631610 | Financial stress (N=776) | infancy (age 8 mo.) | 0.945 | 0.913 | 0.027 | 1.04E-08* | -0.027 | 0.0056 | -0.038 | -0.016 | 19 | 11322739 | DOCK6 | 0 |
| cg06783003 |  | infancy (age 1.75) | 0.856 | 0.894 | 0.024 | 5.42E-08 | 0.036 | 0.0083 | 0.020 | 0.052 | 1 | 45116008 | RNF220 | 0 |
| cg01050704<sup>a</sup> |  | preschool (age 5) | 0.017 | 0.019 | 0.025 | 5.67E-08 | 0.002 | 0.0005 | 0.001 | 0.003 | 19 | 59084995 | MZF1-AS1 | 0 |
| cg02006977 |  | preschool (age 5) | 0.015 | 0.017 | 0.024 | 8.21E-08 | 0.002 | 0.0005 | 0.001 | 0.003 | 12 | 69139955 | SLC35E3 | 0 |
| cg21299458 |  | preschool (age 5) | 0.113 | 0.155 | 0.035 | 2.54E-11* | 0.038 | 0.0070 | 0.024 | 0.051 | 22 | 20779896 | SCARF2 | 0 |
| cg19219503 |  | middle childhood (age 7) | 0.917 | 0.869 | 0.030 | 2.77E-10* | -0.034 | 0.0070 | -0.048 | -0.020 | 10 | 37414802 | ANKRD30A | 0 |
| cg21924472 |  | recency | 0.755 | 0.772 | 0.027 | 1.86E-08 | 0.003 | 0.0006 | 0.002 | 0.004 | 4 | 139600734 | LINC00499 | 255235 |
| cg24996440 |  | recency | 0.567 | 0.587 | 0.026 | 2.88E-08 | 0.004 | 0.0009 | 0.003 | 0.006 | 2 | 3583570 | RNASEH1 | 9119 |
| cg00928478 | Neighborhood disadvantage (N=704) | infancy (age 1.75) | 0.021 | 0.018 | 0.027 | 2.68E-08 | -0.002 | 0.0005 | -0.003 | -0.001 | 10 | 99078824 | FRAT1 | 196 |
| cg01954337 |  | infancy (age 1.75) | 0.051 | 0.061 | 0.028 | 6.29E-08 | 0.008 | 0.0018 | 0.005 | 0.012 | 11 | 3819010 | NUP98 | 0 |
| cg04996689 |  | infancy (age 1.75) | 0.029 | 0.036 | 0.028 | 3.08E-08 | 0.006 | 0.0011 | 0.003 | 0.008 | 5 | 52285560 | ITGA2 | 0 |
| cg12069925 |  | infancy (age 1.75) | 0.042 | 0.052 | 0.029 | 5.40E-09* | 0.007 | 0.0014 | 0.004 | 0.009 | 17 | 11900858 | ZNF18 | 72 |
| cg14522055 |  | infancy (age 1.75) | 0.031 | 0.037 | 0.027 | 7.38E-08 | 0.005 | 0.0011 | 0.003 | 0.007 | 15 | 64338757 | DAPK2 | 235 |
| cg19157140 |  | infancy (age 1.75) | 0.014 | 0.016 | 0.037 | 3.58E-11* | 0.002 | 0.0004 | 0.001 | 0.003 | 7 | 766323 | PRKAR1B | 0 |
| cg21740964 |  | infancy (age 1.75) | 0.160 | 0.177 | 0.025 | 6.62E-08 | 0.013 | 0.0028 | 0.008 | 0.019 | 6 | 3849331 | FAM50B | 299 |
| cg24826892<sup>a</sup> |  | infancy (age 1.75) | 0.016 | 0.019 | 0.030 | 6.03E-09* | 0.003 | 0.0006 | 0.002 | 0.004 | 11 | 71159390 | DHCR7 | 0 |
| cg08546016 |  | preschool (age 5) | 0.051 | 0.057 | 0.029 | 3.93E-09* | 0.006 | 0.0012 | 0.004 | 0.009 | 17 | 72776238 | TMEM104 | 0 |
| cg12412390 |  | middle childhood (age 7) | 0.039 | 0.047 | 0.030 | 9.07E-08 | 0.008 | 0.0016 | 0.005 | 0.011 | 4 | 96469286 | UNC5C | 0 |

DNAm = unadjusted DNA methylation (beta values) averaged within group; Increase in $R^2$ = increase in $R^2$ explained by first hypothesis chosen after accounting for covariates; P = p-value of covariance test assessing significance of increase in $R^2$ explained; Beta, SE, Lower 95% CI, Upper 95% CI = parameter estimate, standard error, and lower and upper limits of 95% confidence interval of regression coefficient of first hypothesis chosen; Chr, Coordinate = chromosome and position of CpG site; Nearest gene = Gene symbol of and distance in bases to nearest gene from CpG site (as measured from transcription start site)
<sup>a</sup>In potentially noisy probe list of Naeem et al. 2014 (i.e., cross-reactive probes, probes with SNPs/INDELs/repeat regions, probes affected by unknown factors) <sup>b</sup>In potentially noisy probe list of Chen et al. 2013 (i.e., cross-reactive probes, probes with SNPs)
\*significant at $p < 1.43 \times 10^{-8}$ , a more stringent p-value threshold that accounted for the testing of seven types of adversity ( $1 \times 10^{-7} / 7 = 1.43 \times 10^{-8}$ )

Across all 40 top sites, exposure to adversity was typically associated with hypermethylation (67.5% positive beta coefficients; *χ*^2^=4.9, p=0.027; **Table 1**). However, exposure to maternal psychopathology and family instability were primarily associated with hypomethylation (6 out of 7 negative beta coefficients, across both types). On average, exposure to adversity during a sensitive period was associated with a 2.4% difference in methylation level (beta) after controlling for all covariates (range 0.1-14.3%). For the two CpG sites associated with recency of exposure to financial stress, one additional adverse event was associated with a 0.3-0.4% increase in methylation per year of age at the event. Of these 40 CpG sites, 17 remained statistically significant after imposing a more stringent p-value threshold that accounted for the testing of seven types of adversity (p=1×10^−7^ / 7=1.43=10^−8^; **Table 1**).

After relaxing the multiple testing correction threshold to a FDR q<0.05, there were 365 CpG sites affected by exposure to adversity (**Figure 3b**; **Table S3**). As with the top 40 Bonferroni-significant sites, methylation at 342 of the 365 FDR-significant sites was best explained by sensitive period models (**Figures 3b**, **Table S3**). Exposure in infancy explained methylation variation at more CpG sites than expected from the background for family instability and neighborhood disadvantage (**Figures S2**). The effects of adversity type and timing on methylation were distributed throughout the genome (**Figure S3**).

### Exposed vs. Unexposed Analysis

Across the seven EWASs, which separately evaluated the effect of ever versus never exposed to each type of adversity on CpG site DNAm, only one statistically significant result emerged (**Figure S4**); this was for cg02431672, a locus located on chromosome 1 79kb away from the gene *FAM183A* and was associated with exposure to abuse (β=−0.005; p=1.58×10^−8^).

Overall, there was very little overlap in identified CpG sites across the top SLCMA and
EWAS results. Most of the top 40 sites had effect estimates that were larger in the SLCMA compared to the EWAS (**Figure 4**). There was also little overlap in findings across specific CpG sites. For example, the cg02431672 locus, which was the top hit in the EWAS of abuse, did not emerge as a top hit in the SLCMA of abuse, failing to appear in the list of FDR significant loci (p=0.0125). Similarly, the top CpG site in the SLCMA (cg21299458), which suggested a sensitive period at age 5 associated with the effects of financial stress, was non-significant in the EWAS of financial stress (β=0.012; p=0.0181; **Figure 5**). These results suggest that the SLCMA allowed us to more effectively identify methylation differences among children with and without a history of exposure to adversity.

**Figure 4.**
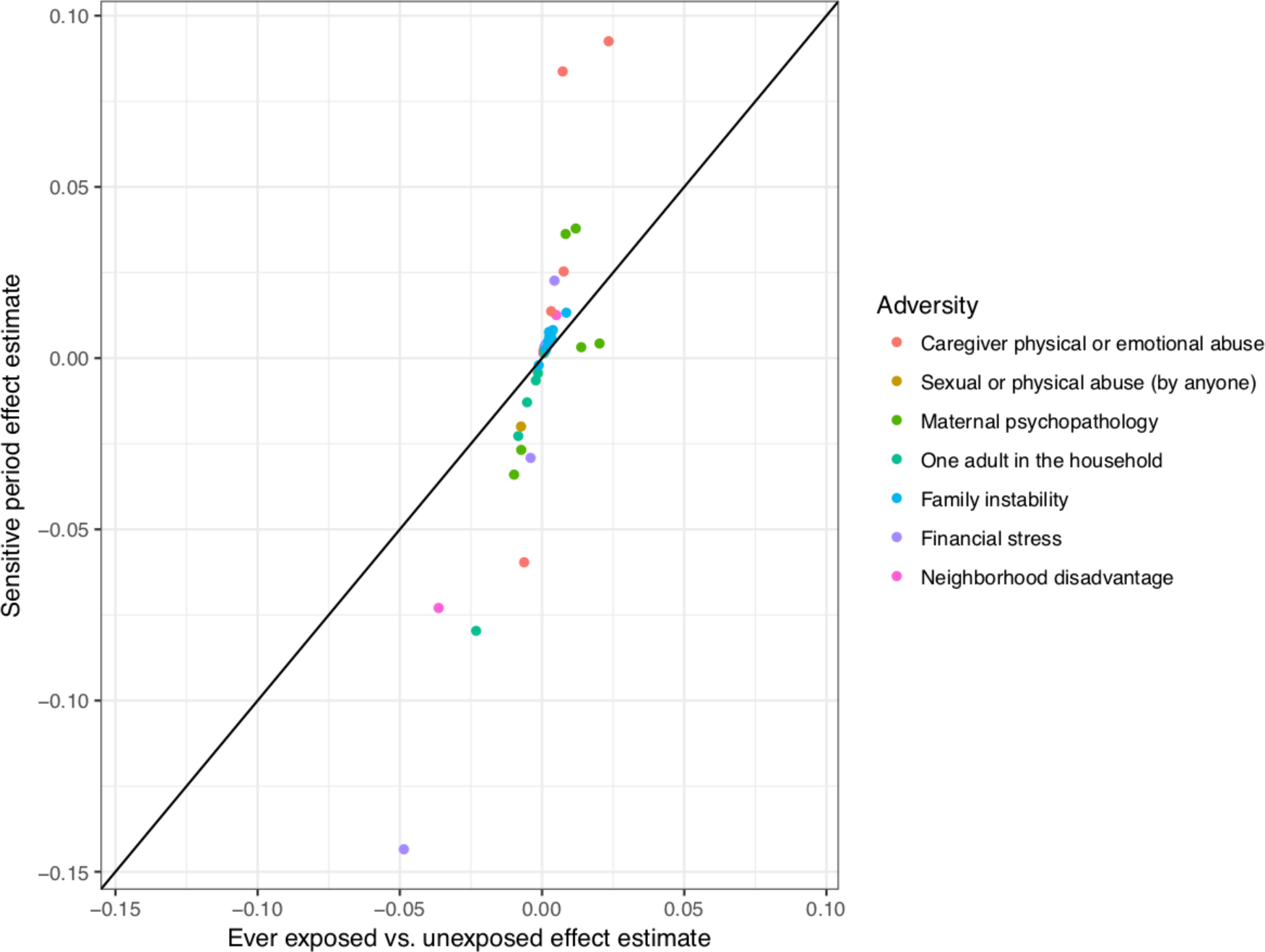
Scatterplot displaying increased power in the SLCMA shown by the comparison of beta estimates from the EWAS vs. SLCMA approaches. In this scatterplot, the y-axis represents the beta estimates associated with the 40 top CpG sites derived for the SLCMA; the x-axis represents the beta estimates associated with the same 40 CpG sites obtained from EWAS. Different types of adversity are indicated by colors. The black straight line denotes the 1:1 correspondence between the two sets of beta values. Substantial deviation from the line suggests increased power in the SLCMA. For most CpG sites, the magnitudes of effect were larger for the SLCMA compared to the EWAS results.

**Figure 5.**
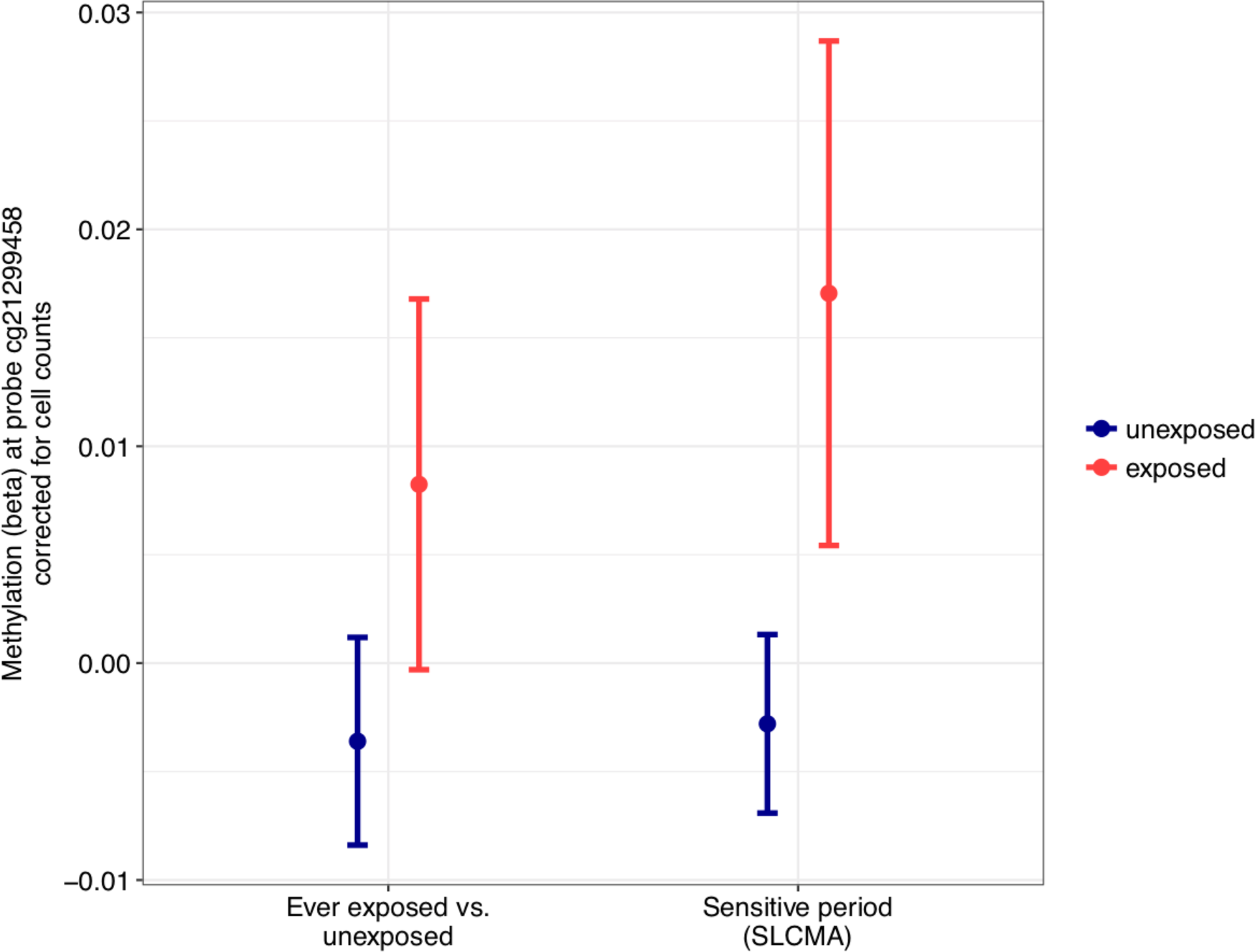
Comparison of EWAS vs. SLCMA estimates for the top CpG site identified in SLCMA, cg21299458. The effect estimates and the confidence intervals obtained from the EWAS approach comparing ever exposed to never exposed to financial stress for cg21299458 are presented on the left. The stage 2 effect estimates and confidence intervals obtained from the SLCMA comparing being exposed to financial stress at age 5 to being unexposed at age 5 for the same CpG site are displayed on the right. The top CpG site in the SLCMA, which suggested a sensitive period at age 5 associated with the effects of financial stress, was non-significant after correction for multiple testing (p=0.0125) in the EWAS of financial stress.

### Sensitivity Analyses

#### Evaluation of the LARS Selection Procedure

There was no evidence in support of compound theoretical models, whereby more than one theoretical model explained the most outcome variability. For each of the top 40 CpG sites, the marginal increase in variance of methylation explained by additional steps of the LARS procedure was not significant (each p>0.05, **Figure S5**), suggesting that methylation was best explained by a single theoretical model.

#### Evaluation of Methylation at Birth for Top CpG Sites

Adversity-associated methylation differences occurred during early childhood for most top CpG sites. For all but two sites, the age 7 DNAm differences between exposure groups were not present at birth (p>0.05/40=0.00125), though the direction of DNAm differences were similar between birth and the other time points for many sites (**Table S4**). An example of a site differentially methylated at birth and an example of a site non-differentially methylated at birth are shown in **Figure S6**.

#### Correction for Genetic Variation

Genetic variation did not appear to influence observed DNAm differences at the top CpG sites. Using a database of methylation quantitative trait loci (mQTLs) of the ARIES cohort (78), there were 627 SNPs associated with DNAm at 17 of the top 40 sites. After controlling for genetic variation at mQTLs linked to these 17 sites, the effect of exposure to adversity remained significant (each FDR q<0.05; **Table S5**), suggesting that adversity could have caused these methylation differences distinct from genetic sequence variation.

### Exploring the Biological Significance of Findings

#### Correlation Between Blood and Brain Tissue

On average, methylation in blood at the top 40 sites was slightly positively correlated with methylation in four brain regions (prefrontal cortex: r_avg_=0.12, entorhinal cortex: r_avg_=0.16, superior temporal gyrus: r_avg_=0.14, cerebellum: r_avg_=0.08; **Table S6**). CpG sites with methylation that is highly correlated between blood and brain tissue may be indicative of important inter-individual covariation (i.e., because of adversity) or a strong genetic influence on methylation, while those that are uncorrelated may still be biomarkers of a response to adversity.

#### Enrichment of Regulatory Elements

As compared to all autosomal loci tested, FDR-significant loci were more likely to be located in gene promoters (*χ*^2^=13.02, p<0.0005) and less likely to be in gene enhancers (χ^2^=3.90, p=0.048; **Figure S7A**). Furthermore, the location of FDR-significant loci differed from all other loci tested relative to CpG Islands (*χ*^2^=36.48, p<0.0001; **Figure S7B**). With eFORGE 1.2 (79), we also tested whether FDR-significant loci colocalize with markers of transcriptional activity. FDR-significant loci were not enriched for DNase I hypersensitivity sites or histone marks in any tissue or cell-type after correction for multiple comparisons (each q>0.05). The strongest trend for enrichment was detected in the analysis of all histone marks in derived neuronal progenitor cells (uncorrected p=0.0003). Annotations at each FDR-significant site are presented in **Table S3**.

#### Biological Processes Potentially Affected by Adversity

Genes near the FDR-significant sites (n=354 genes) corresponded to 158 clusters of GO biological process terms (75). The top 8 GO term clusters, including circulatory system development, cell proliferation and migration, and neuronal development, were more likely to be represented than chance (average enrichment p<0.05; the top 15 clusters are presented in **Figure S8**).

Additionally, we uncovered evidence of functional constraint for these genes. Genes annotated to FDR-significant sites were more highly constrained, as measured by the probability of intolerance to Loss-of-Function variation (pLI) from ExAC (76), than the rest of the autosomal genes tested (permutation p=0.0001; **Figure S9**). This indicates a greater importance of these genes, on average, to survival and reproduction over human evolution.

#### Phenotypic Associations for Genes Near Top Sites

We searched the NHGRI-EBI GWAS Catalog (GRASP) (77) to identify whether common variants in genes corresponding to our 40 top CpG sites were associated with relevant phenotypes. Six genes mapped to our top sites had SNPs associated with psychiatric or neurological phenotypes at a genome-wide suggestive level (p<1×10^−5^; **Table S7**), underscoring the possible biological significance of these methylation differences.

## Discussion

This prospective study used data from a large population-based sample of children to test three competing life course theoretical models describing the association between exposure to childhood adversity, measured repeatedly across the first 7 years of life, and DNAm at age 7. By comparing these theoretical models to each other, we could evaluate which one explained the most variation in DNAm. To our knowledge, this is the first use of the SLCMA in an epigenome-wide context.

The main finding of this study is that the effect of adversity on DNAm depends primarily on the developmental timing of exposure. In our Bonferroni-corrected analysis, we identified 40 CpG sites that were differently methylated following exposure to adversity, with more than half of these loci showing associations based on adversity occurring during infancy, meaning before age 3. These results are consistent with at least one human longitudinal study (16) and multiple animal studies (21, 22, 24, 25) in emphasizing the existence of sensitive periods (19, 20) - particularly occurring shortly after birth - when epigenetic programming is maximally dynamic in response to parental care disruptions and other environmental inputs. The lack of detectable sensitive periods in one recent study (32) may be due to focusing only on adversities occurring at or after 5 years of age. Interestingly, neither the accumulation nor recency of the adversity explained considerable variability in DNAm. The observed DNAm differences were absent at birth, identified for a range of adversities, and unrelated to genetic variation. The absence of support for an accumulation model is surprising, given previous research linking cumulative time spent in institutional care to DNAm status in key stress-related genes (29).

Perhaps more importantly, our results suggest that broad classifications of individuals as exposed versus unexposed to “early life” adversity - although commonly used - may dilute observed effects and fail to detect DNAm differences among those exposed to adversity during specific life stages. The lack of overlap in identified loci across the SLCMA and EWAS suggest that refinement of the environmental phenotype - by treating each time point of exposure (and its accumulation) as unique - may better capture underlying signal. Indeed, results of a post-hoc power calculation suggest that the EWAS of exposed versus unexposed will be underpowered when the true underlying relationship between exposure and outcome depends on the timing or amount of exposure (**Supplemental Materials**).

Although these findings emphasize the importance of exposure timing, greater insights are needed regarding the age stages when adversity may be most harmful, as mixed results have emerged among the small number of studies comparing the effects of “early” to “later” adversity. Some retrospective studies have shown that adolescent DNAm patterns are more strongly associated with exposure to life stress during adolescence more than with earlier adversity (27). However, other studies have found potentially persistent effects of childhood adversity into adolescence (80) and adulthood (81), even after accounting for subsequent stress exposure. A recent study also found that the effects of the timing of adversity may be gene-specific (29). As epigenetic patterns appear to vary over the life course (26, 82), longitudinal studies are needed to study the developmental trajectories of DNAm and evaluate the extent to which these adversity-induced DNAm differences persist or attenuate over time, and operate independently of or in interaction with subsequent experience to ultimately predict disease outcomes.

Several limitations are noted. First, some adversity measures were drawn from single items. Parents may have also under-reported exposure to stigmatizing experiences (83, 84), especially if they were implicated in the exposure (85). However, the prevalence of several adversities, including those capturing possible experiences of child abuse, were similar to and even greater than those reported from some nationally-representative samples (9, 86). Second, as with any longitudinal study, there was attrition over time, which could result in bias due to loss of follow-up. However, ARIES children were sampled from among those with the most complete longitudinal data. Within the field of epigenetics, efforts are now underway to understand the consequences of attrition and how potential biases arising from attrition could be mitigated through multiple imputation or other strategies. Third, we were unable to examine the impact of experiencing multiple adversities simultaneously because these adversities were measured at different time points. Fourth, the DNAm samples were obtained from peripheral tissue and not the brain; multiple datasets, however, are starting to identify limited though important shared DNAm patterns across central nervous system and peripheral tissue (87). Finally, the p-values derived from the covariance tests could be potentially inflated, as the test relies on asymptotic theories and therefore does not theoretically guarantee the control of Type I error rate in a finite sample (70). However, the covariance test might be a more sensitive method to detect signals compared to other post-selection significance tests that make fewer assumptions (88). As the relative statistical power of the available tests remains unclear, future simulation studies are much needed to identify the best inference tools in different settings.

In summary, this study lends further support to the evidence base showing that DNAm patterns are responsive to experience. However, these results reveal that DNAm patterns may be most influenced by exposures during sensitive periods in development. Efforts may therefore be needed to move beyond crude comparisons of those exposed versus unexposed to “early life” adversity.

## Acknowledgments

We are extremely grateful to all the families who took part in this study, the midwives for their help in recruiting them, and the whole ALSPAC team, which includes interviewers, computer and laboratory technicians, clerical workers, research scientists, volunteers, managers, receptionists and nurses. The UK Medical Research Council and the Wellcome Trust (Grant ref: 102215/2/13/2) and the University of Bristol provide core support for ALSPAC. ARIES was funded by the BBSRC (BBI025751/1 and BB/I025263/1). Supplementary funding to generate DNA methylation data which is included in ARIES has been obtained from the MRC, ESRC, NIH and other sources. ARIES is maintained under the auspices of the MRC Integrative Epidemiology Unit at the University of Bristol (MC_UU_12013/2 and MC_UU_12013/8). This publication is the work of the authors, each of whom serve as guarantors for the contents of this paper. This work was conducted with support from Harvard Catalyst | The Harvard Clinical and Translational Science Center (National Center for Research Resources and the National Center for Advancing Translational Sciences, National Institutes of Health Award UL1 TR001102 and K01MH102403) and financial contributions from Harvard University and its affiliated academic healthcare centers. The content is solely the responsibility of the authors and does not necessarily represent the official views of Harvard Catalyst, Harvard University and its affiliated academic healthcare centers, or the National Institutes of Health. The authors thank Kathryn Davis, and Samantha Ernst for their assistance in preparing this manuscript for publication.

